# TnFLX: a third-generation mariner-based transposon system for *Bacillus subtilis*

**DOI:** 10.1101/825950

**Authors:** Felix Dempwolff, Sandra Sanchez, Daniel B. Kearns

## Abstract

Random transposon mutagenesis is a powerful genetic tool to answer fundamental biological questions in an unbiased approach. Here, we introduce an improved *mariner-based* transposon system with higher stability, and with versatile applications. We take advantage of the lower frequency of unintended recombination during vector construction and propagation in a low copy number system in *E. coli* to improve construct integrity. We generated a variety of transposons allowing for gene disruption or artificial overexpression each in combination with one of four different antibiotic resistance markers. In addition, we provide transposons that will report gene/protein expression due to transcriptional or translational coupling. We believe that the TnFLX system will help enhance flexibility of future transposon modification and application in *Bacillus* and other organisms.

**Importance:** The optimization of transposase encoding vectors in terms of stability during cloning and propagation is crucial for the reliable application of this system in any host organism. With an increased number of antibiotic resistance markers and the possibility to detect translational activity, the TnFLX transposon system will significantly help the implication of forward genetic methods in the field of cellular biology.

## INTRODUCTION

Transposon mutagenesis is a powerful method in molecular genetics, as it combines random insertional mutagenesis with the generation of linked antibiotic markers and rapid insertion site identification. Typically, an antibiotic resistance cassette is cloned between two inverted terminal repeat elements (ITR elements) that define the transposon. Next, the transposase enzyme is provided as purified protein *in vitro* or expressed from a gene *in vivo* to recognize and mobilize the transposon to another random genetic location (1–3). Different transposon systems have different transposition frequencies, different insertion biases, and different mechanisms for providing the transposase. Moreover, the functionality of transposons can be extended by cloning additional genetic material between the IS elements. For instance, transposon derivatives have been generated that insert not only the antibiotic resistance cassette but other genes and promoters for the generation of randomly inserted reporter fusions and artificial expression systems (4–8).

*Bacillus subtilis* is a model genetic organism and a number of transposon systems have been developed to aid genetic analyses. The earliest transposon system developed for *B. subtilis* was *Tn917* which generated random insertions but exhibited a strong bias toward insertions near the chromosome’s terminus (9–11). The next system used transposon Tn*10* that improved insertion diversity and provided an easier method for insertion site identification but was still prone to insertions at particular locations or “hotspots” (12–16). Later the first generation of systems using transposon *mariner* was developed that inserted at two-base pair “TA” sequences in the chromosome thus increasing the potential number of insertion sites in low G+C content organisms (2). Sequencing of transposon pools (TnSeq) indicates that mariner-based transposition is highly random and seems to alleviate “hotspotting” issues of the Tn*10* and Tn*917*-based systems (17–19). Later, our group constructed second generation derivatives of the *mariner* system adding additional antibiotic cassettes and transposons for random transcriptional reporter insertions of the β-galactosidase gene *lacZ* and random artificial expression insertions with an outward facing IPTG-inducible promoter (20).

After sharing *E. coli* strains carrying the *B. subtilis* transposon delivery plasmids with other labs, it became clear that there was a problem with our system. Labs were having difficulty recovering intact plasmids from *E. coli* and isolation of plasmid DNA from four different colonies of the same frozen *E. coli* strain gave four different plasmid digestion patterns, none of which were correct (Fig 1A). We retracted the plasmids from the Bacillus Genetic Stock Center (BGSC, Ohio State University) and provided alternative solutions for obtaining the transposon system we created (retraction letter in supplemental material).

**Figure 1:**
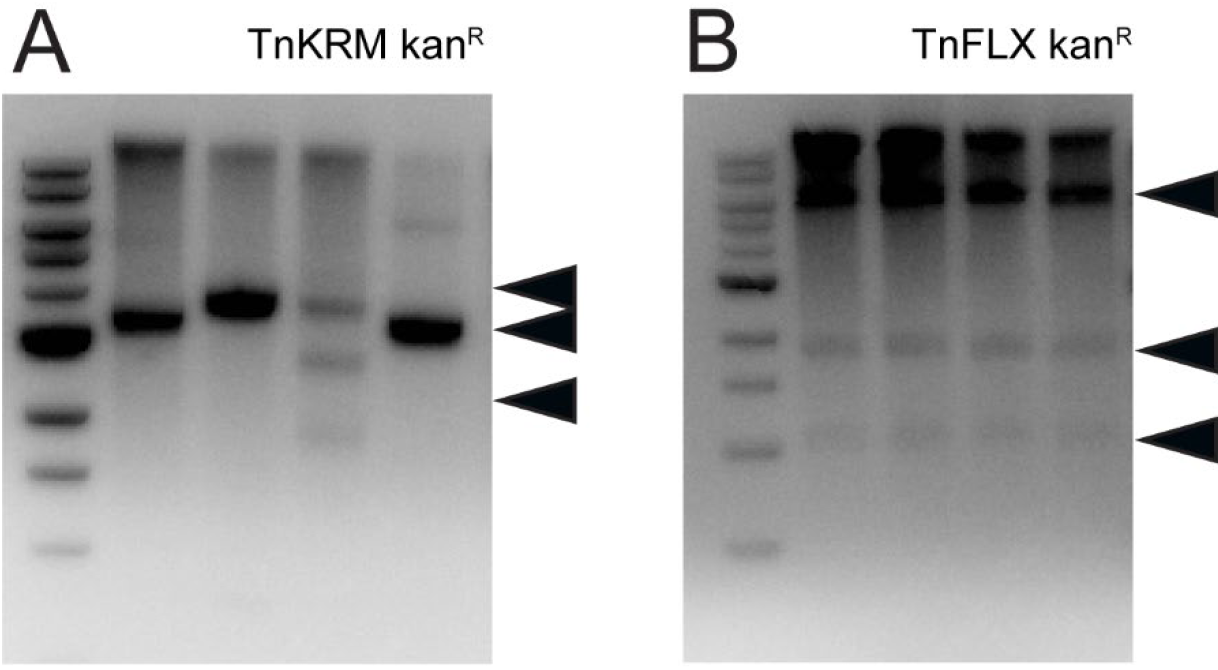
Plasmid integrity of the TnKRM (pKB178) and the TnFLX (pFK131) series. Analysis of a restriction digests of individually isolated plasmids from clonal cultures. A pTnKRMKan was incubated with *NheI* and *Eco*RI. Fragment sizes of approx. 3900, 2850, and 1450 bp are expected. B pTnFLXABKan was incubated with *SalI*. Fragments sizes of approx. 6600, 1950, and 1100 bp are expected. The 1 kb DNA ladder (New England biolabs) was used for size determination. Arrowheads indicate expected fragment sizes.

Here we report the construction of a third-generation of *mariner-based* transposons that affords increased stability in *E. coli* through the use of a low-copy number delivery plasmid. Moreover, we regenerate all of the second-generation plasmids using the new platform, add an additional antibiotic resistance cassette conferring resistance to tetracycline, and create a new transposon derivative for the generation of random C-terminal translational fusions to green fluorescent protein (GFP).

## RESULTS AND DISCUSSION

### Redesign of the transposon delivery system

*Himar-based* systems for random transposon mutagenesis were already described for *B. subtilis* ((21)(20)). Unfortunately, while working with the existing TnKRM series of *himar* based transposon system we realized a high frequency of genetic reorganization in the delivery plasmid during cloning and proliferation in an *E. coli* DH5α host. Purification of plasmids from separate clonal isolates of *E. coli* indicated a heterogeneous pattern of fragments, none of which resembled the digestion pattern predicted for the intact plasmid (Fig. 1A). We attribute this inconsistency of the genetic arrangement to recombination events performed by the plasmid-encoded transposase. Transposase expression might be high in *E. coli* as the *himar* gene was under the conserved and constitutive σ^70^/σ^A^ promoter sequence of *E. coli/B. subtilis*. Moreover, the replication origin of the transposon delivery plasmid was derived from high-copy number pUC plasmids (22), further amplifying the potential for *himar* expression. We infer that expression of the *himar* transposase likely contributes to high-frequency transposition, plasmid rearrangements, and a loss of plasmid sequence integrity including the loss of the transposon itself.

In an effort to increase plasmid fidelity, we chose to design a new transposon delivery system, based on a low copy number plasmid and a modified promoter for the transposase encoding gene. As the vector backbone, we chose the pGB2 (23) plasmid (kind gift of T. Bernhard, Harvard University). It consists of a low activity origin of replication (ori_*Ec*_) that will only allow for a few replication cycles during the generation time of a *E. coli* cell and a spectinomycin resistance cassette (spec_*Ec*_^R^) for plasmid maintenance during passage in a *E. coli* host. An erythromycin resistance cassette (erm^R^) was integrated into pGB2 to allow for selection of plasmid uptake in the *B. subtilis* host. Next, we integrated the *himar* expression construct of the plasmid pMarB (21), so transposase expression was under the control of a *B. subtlis* promoter recognized by the stress response sigma factor σ^B^, further reducing the likelihood of expression in *E. coli*. To complete our transposon delivery system (pFK7) (Fig 2), we integrated a temperature sensitive origin of replication specific for *B. subtilis* (ori*Bs*^Ts^) to allow replication in the host at permissive temperatures (<30°C) (24–27). In addition, we included a single intergenic *SmaI* restriction site for blunt end linearization, allowing for the integration of various transposable elements into this system by isothermal assembly (28).

**Figure 2.**
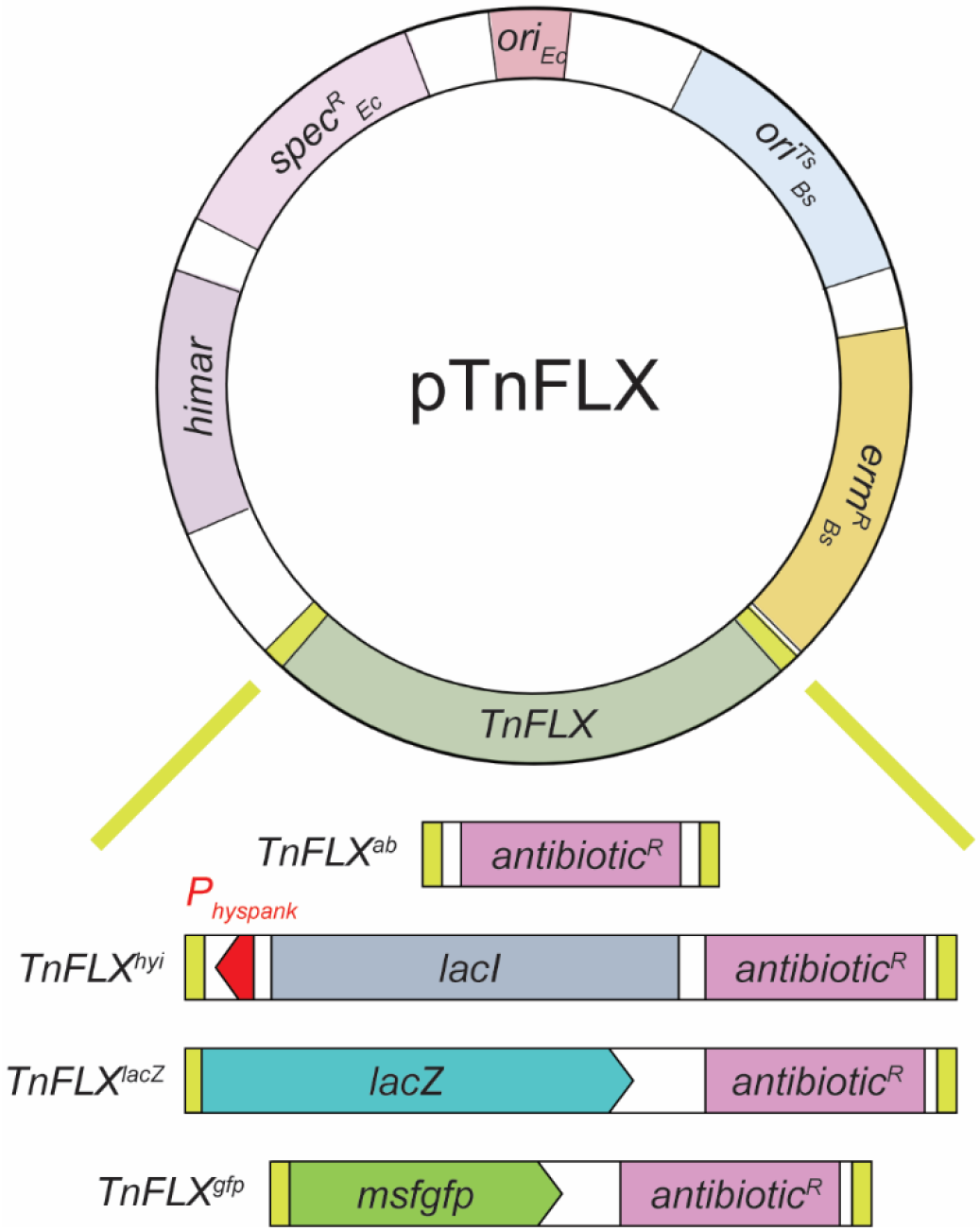
The pTnFLX delivery plasmid. Schematic representation of the TnFLX series of plasmids. ori_Ec_ origin of replication (*E. coli)*, Spec^R^_Ec_ spectinomycin resistance cassette, himar, σ^B^ controlled mariner transposase, ori^Ts^_Bs_ temperature sensitive origin of replication (*B. subtilis*), erm^R^ erythromycin resistance cassette, TnFLX, inverted terminal repeats flanking the transposable region. *Sma*I, endonuclease recognition site. TnFLX transposons introduced in this study. Flanking yellow rectangles represent inverted terminal repeats. Red arrow indicates promoter orientation.

### Construction of TnFLX and reconstruction of the TnKRM series of transposons

The first generation of *mariner-based* transposon system included a *mariner* transposon containing a single antibiotic resistance cassette conferring kanamycin resistance (TnYLB-1) (2). The second generation of modified *mariner* transposons, TnKRM, introduced transposons that had the kanamycin resistance cassette as well as two other transposons separately carrying resistance to spectinomycin and chloramphenicol respectively (20). Here we constructed transposons separately carrying genes conferring resistance to kanamycin (kan^R^), spectinomycin (spec_Bs_^R^), chloramphenicol (cat^R^) and a fourth cassette conferring resistance to tetracycline (tet^R^) (Fig. 2). In addition, all antibiotic resistance cassettes were flanked by a common short nucleotide sequence, which will allow for transposon insertion-site identification with the same pair of oligonucleotides (with the exception of the TnFLXPhyi series that requires a distinct set of oligonucleotides) (see table 1).

**Table 1.**
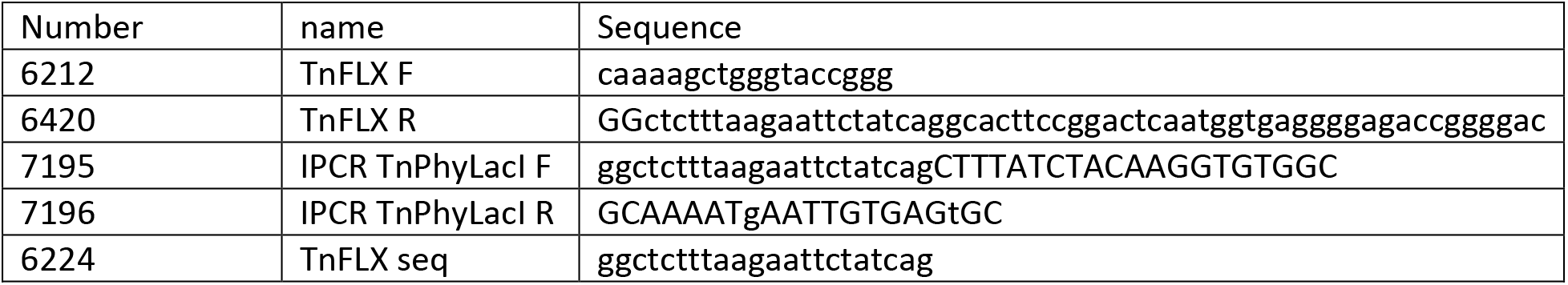
oligonucleotides used for the identification of the transposon insertion site

As proof-of-concept each transposon was introduced into *B. subtilis* NCIB3610 cells (29). Cells were propagated in LB medium at room temperature overnight, diluted and regrown in LB medium at the non-permissive temperature of 42° C for 6 hours, dilution plated on LB-agar plates containing the transposon specific antibiotic and incubated overnight at 42° C. Whereas wild type colonies have a dull, rough architecture, mutant colonies were isolated that exhibited a shiny, smooth colony morphology indicative of a biofilm defect (30). Inverse PCR was used to identify the location of each transposon insertion and each strain was found to be disrupted in a gene previously reported to be involved in biofilm formation (Table 2). One mutant was isolated with a hyper rough phenotype indicative of EPS overproduction and one mutant was isolated with a mucoid colony morphology indicative of poly-γ-glutamate overproduction. Again, each mutant was disrupted for a gene previously reported to confer the associated phenotype (Table 2). We conclude that the TnFLX series of transposons randomly insert in the chromosome and create loss-of-function mutations.

**Table 2.**
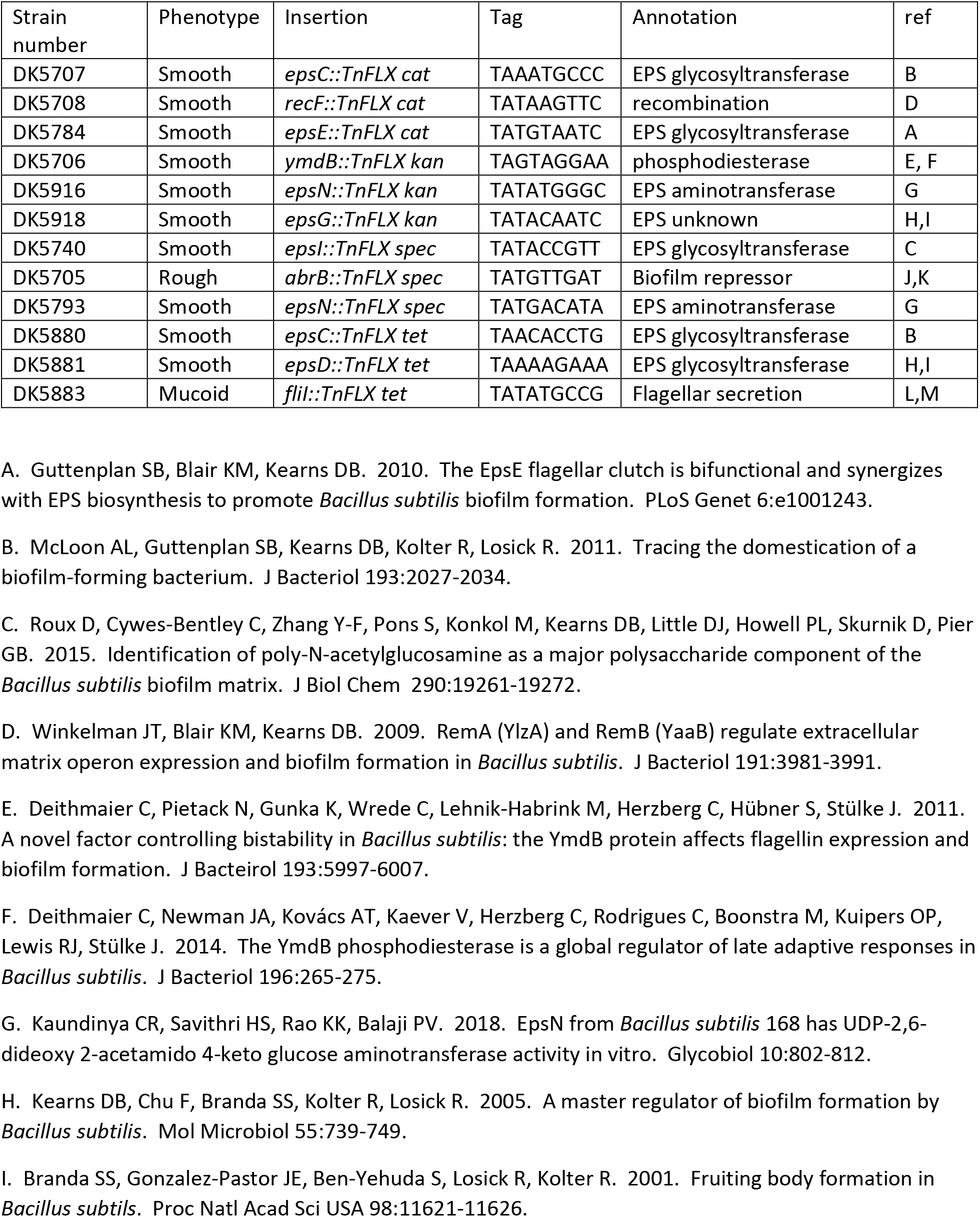
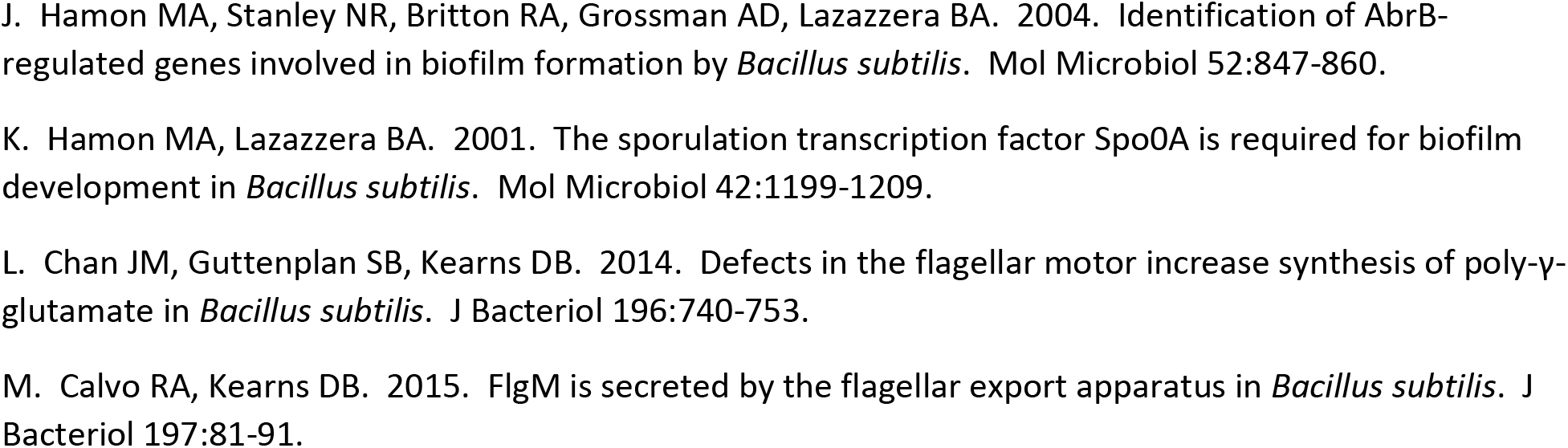
isolated TnFLX mutants

In addition to gene disruption, the artificial expression of a gene in a non-physiological context can reveal valuable information about the function of a gene product. To allow for this kind of artificial control over gene expression, a transposon TnHyJump was created with an antibiotic resistance cassette and a strong P_hyspank_ promoter (31) cloned facing outward from one arm of the transposon. The *lacI* gene encoding the P_hyspank_ repressor protein LacI could be separately integrated at an ectopic location in the chromosome to provide IPTG-inducible control. Here, we created the TnFLXPhyi series of transposons in which an antibiotic resistance cassette was also combined with the outward facing P_hyspank_ promotor (kind gift from D. Rudner, Harvard University). In the TnFLXPhyi case, however, the *lacI* gene was also encoded between the insertion elements such that IPTG-inducible control was inherent with, and linked to, each insertion (Fig. 2).

The TnFLXPhyi system was validated by performing a transposon mutagenesis in *B. subtilis* PY79 cells. Overexpression blue/white screens using the previous generation TnHyJump system indicated that insertions upstream of cryptic β-galactosidase genes could confer a blue color (20). Therefore, after mutagenesis, cells were spread on plates containing the transposon-specific antibiotic, the inducer IPTG and the indicator X-Gal, and incubated overnight at the non-permissive temperature. Subsequently, plates were examined for blue colonies, indicating the enzymatic cleavage of X-gal. Blue colonies were observed at a low frequency of approximately one per thousand (Fig 3A). We isolated 12 blue colonies (3 for each different transposon used) (see table 3) and identified the transposon insertion site by IPCR (Table 1). Insertions of the transposon in 9 out of 12 mutants were found to be within the large *yes* operon and oriented such that the P_hyspank_ promoter could drive artificial expression of the *yesZ* gene encoding a cryptic β-galactosidase (table 3). For one strain the transposon was integrated such that the P_hyspank_ promoter was inserted in, and co-oriented with, the coding sequence for *ganQ*. Analysis of the downstream region of the *ganQ* encoding operon showed the presence of the gene encoding for another β-galactosidase, GanA (formerly LacA) (32). Finally, two transposons were found to be inserted within the *ganR* gene, and unlike the other 10 insertions, the blue color was IPTG-independent (Fig. 3B). Consistent with IPTG- independence, disruption of *ganR* encoding GanR (formerly LacR) derepressed GanA expression (32). Thus, each TnFLXPhyi transposon insertion was capable of generating insertional disruptions, or artificial expression of downstream adjacent genes.

**Figure 3.**
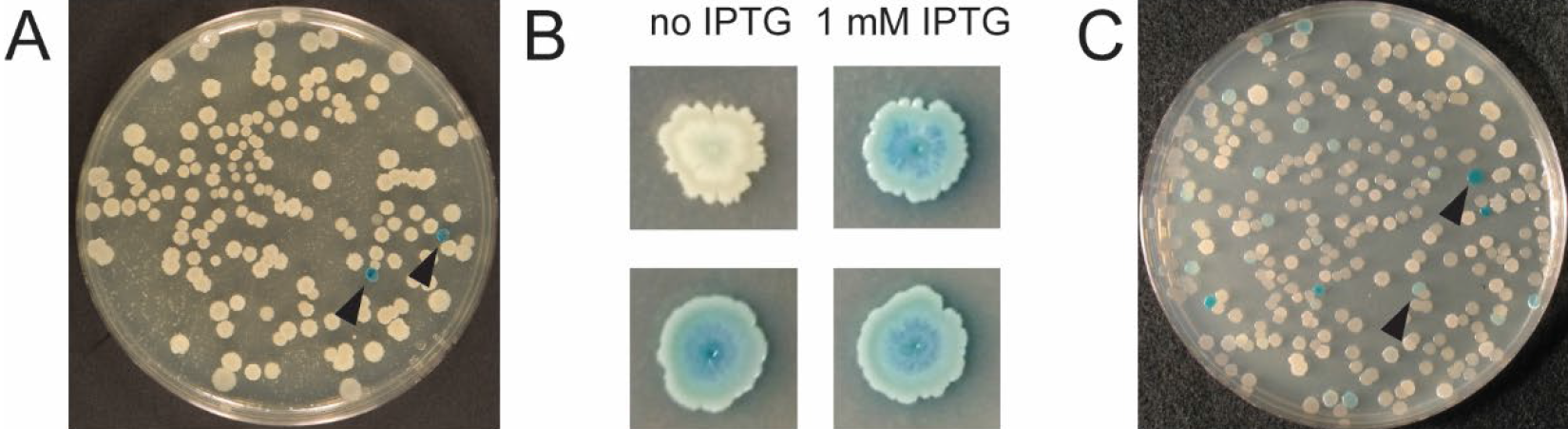
colorimetric analysis of transposon mutagenesis experiments. A TnFLXPhyi series. The occurrence of blue colonies is a rare event (arrow heads indicate blue colonies). B Blue colonies isolated from a TnFLXPhyi screen. Most strains showed inducer responsivity (upper panel). At a low frequency, strains that were unresponsive to IPTG addition were isolated (lower panel). C TnFLXLacZ transposon series. Blue colonies can be observed at a high frequency.

**Table 3.** isolated TnFLXPhyi mutants

| Strain number | Phenotype | Insertion | Tag | IPTG responsive |
| --- | --- | --- | --- | --- |
| DK7507 | Blue on X-Gal plates | *yesX*::TnFLXPhyl cat | TATCATCGG | Yes |
| DK7508 | Blue on X-Gal plates | *yesS*::TnFLXPhyl cat | TATGGCCGC | Yes |
| DK7509 | Blue on X-Gal plates | *yesS*::TnFLXPhyl cat | TATGGAGAG | Yes |
| DK7510 | Blue on X-Gal plates | *ganQ*::TnFLXPhyl spec | TAACCTTTA | Yes |
| DK7525 | Blue on X-Gal plates | *yesW*::TnFLXPhyl spec | TAGTTGCCT | Yes |
| DK7527 | Blue on X-Gal plates | *yesX*::TnFLXPhyl spec | TATGGTGTA | Yes |
| DK7505 | Blue on X-Gal plates | *ganR*:: TnFLXPhyl tet | TATGTGGGC | No |
| DK7692 | Blue on X-Gal plates | *yesQ*::TnFLXPhyl tet | TAATGCTCG | Yes |
| DK7506 | Blue on X-Gal plates | *yesR*:: TnFLXPhyl tet | TACAAAATA | Yes |
| DK7528 | Blue on X-Gal plates | *yesQ*::TnFLXPhyl kan | TATATTATG | Yes |
| DK7530 | Blue on X-Gal plates | *ganR*::TnFLXPhyl kan | TAAGTCATC | No |
| DK7532 | Blue on X-Gal plates | *rhgT*::TnFLXPhyl kan | TACCTTGTT | Yes |

The previous generation of transposons introduced another functional application to the transposon mutagenesis system in which the gene encoding the β-galactosidase LacZ was cloned with an antibiotic resistance cassette within the boundaries of the transposon insertion elements, TnLacJump (20). Thus, insertions of the transposon into, and co-oriented with, an actively transcribed region of the chromosome would create a transcriptional fusion to *lacZ* (in addition to gene disruption depending on the insertion site). To recreate TnLacJump functionality, the *lacZ* gene was cloned into the TnFLX system along with an antibiotic resistance cassette. As proof-of-concept, after mutagenesis, cells were spread on plates containing the transposon specific antibiotic, and the indicator X-Gal and incubated overnight at the non-permissive temperature. Blue colonies were clonally isolated (Fig. 3C), and the insertion site was identified by IPCR. In each case, the transposon had inserted into an open-reading-frame and was cooriented with the direction of transcription consistent with the generation of an inserted transcriptional reporter (table 4). We conclude that the TnFLXlacZ system (Figure 2) generates random lacZ insertions.

**Table 4.**
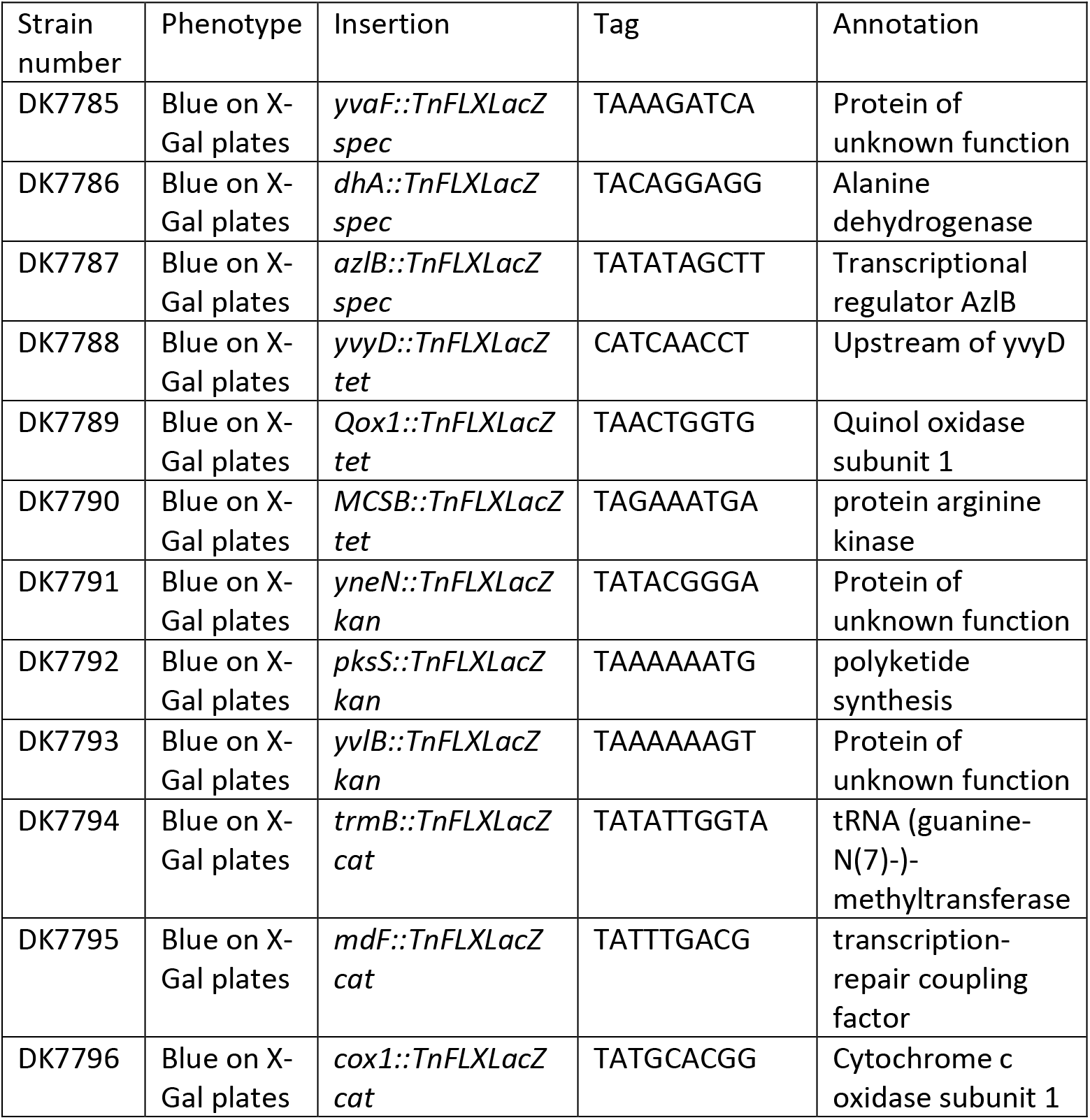
isolated TnFLXLacZ mutants

### TnFLXgfp

Fluorescent translational fusion proteins are a powerful tool to investigate temporal and spatial protein activity, but fluorescent fusions are available for only a subset of the *B. subtilis* proteome. Here, we introduce a new family of transposons, in which a *gfp* gene was cloned into the TnFLX system with an antibiotic resistance cassette (Figure 2). For reasons of high photostability and quantum yield as well of its enhanced folding capabilities, we chose to base the TnFLXgfp series on the monomeric version of the super folder green fluorescent protein (msfGFP) (33, 34). A coding sequence of *msfgfp*, which was optimized for *B. subtilis* codon usage, was combined with an antibiotic cassette and flanked by *himar* specific ITRs. The ITR and *msfgfp* were positioned in a way that no base triplet upstream of the fluorophore-encoding region would lead to premature translational termination. Moreover, the *msfgfp* sequence lacked a Shine-Dalgarno ribosome binding site ensuring that GFP would only be expressed when inserted in-frame to create a C-terminally GFP fusion protein in the protein’s genetic context on the chromosome.

For an initial test, a transposon mutagenesis was performed by selecting for the corresponding antibiotic resistance at the non-permissive temperature, approximately 400 000 colonies were harvested and the combined pool was observed by fluorescence microscopy. A small subfraction of cells in the initial pool (approximately one in a thousand) displayed a detectable msfGFP-specific signal, indicating successful integration into an actively transcribed open reading frame (Figure 4A). To enrich for cells expressing detectable levels of msfGFP, the pool was subjected to fluorescence activated cell sorting (FACS). Cells displaying fluorescence levels higher than the wild type strain were separated, recovered by incubation on LB-agar plates and pooled to generate the msfGFP-positive mutant library (Figure 4A). From these mutant libraries, 12 mutants (3 for each of the 4 transposons) were isolated, the localization pattern was assessed, and the transposon insertion site determined by IPCR.

**Figure 4.**
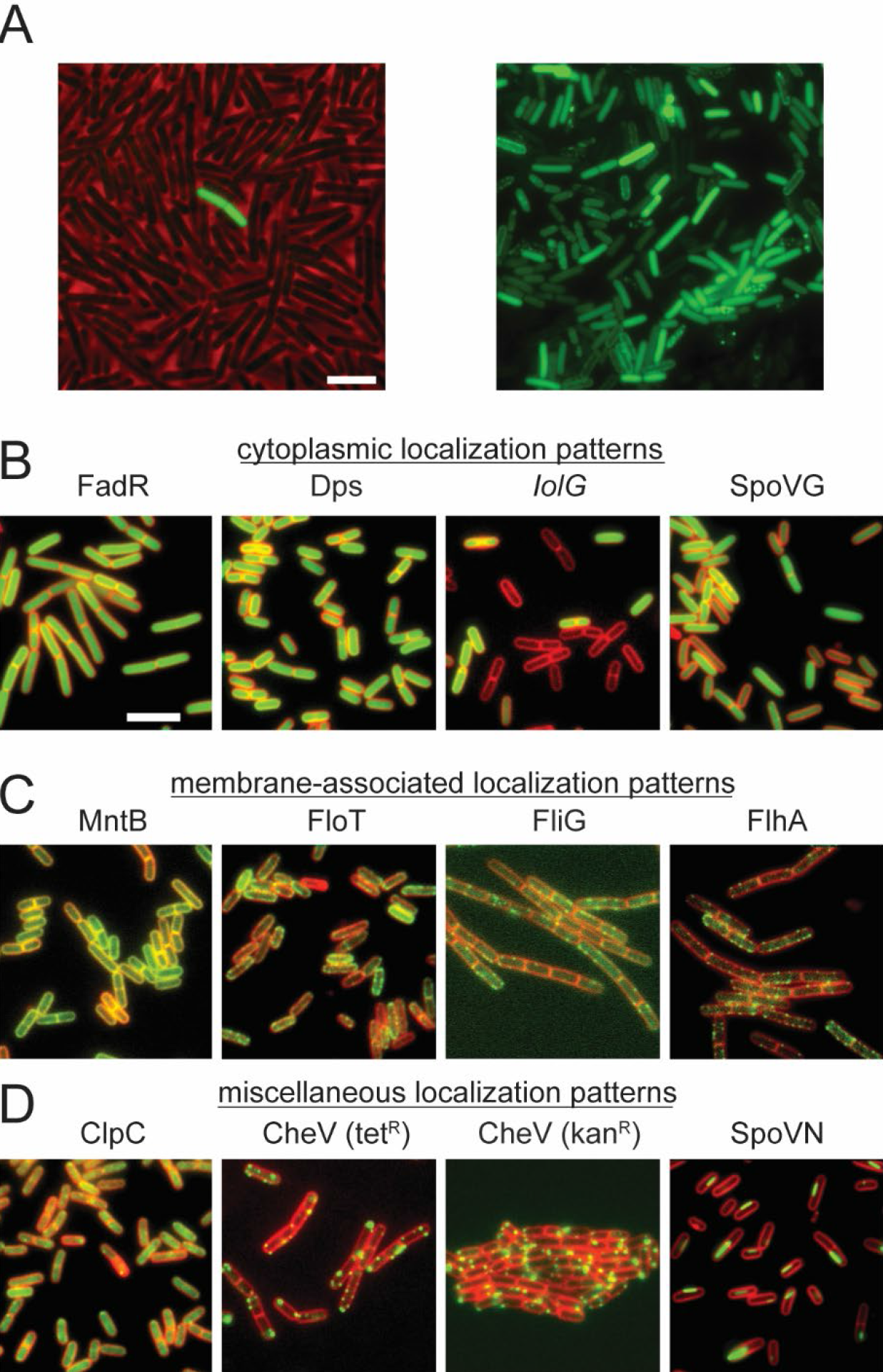
TnFLXGFP analysis. A Analysis of the initial transposon mutant pool by fluorescence microscopy. Before sorting (left,) only one cell of the approx. >100 cells in the field of view displays a detectable GFP signal (image represents an overlay of the phase contrast (red) and the GFP channel (green). After enrichment for cells expressing a functional GFP fusion protein by FACS, a lot of different GFP localization patterns can be observed (right). B-D. Analysis of clonal isolates from the enriched GFP library. GFP localization is shown in green, stained membranes are shown in red. Bars represent 5 μm.

A variety of subcellular localization patterns were observed in the mutants containing fluorescent GFP fusions. One pattern of expression was the diffuse localization of green fluorescence in the cytoplasm such as those in which the transposon inserted into the *fadR* gene, encoding the transcriptional repressor FadR, and the *dps* gene, encoding the iron storage protein Dps (Fig. 4B). Either these proteins normally localize diffusely in the cytoplasm or insertion of the transposon and premature truncation of the coding sequence in favor of the GFP fusion altered localization. At most, we can infer that the expression of these genes occurs during vegetative growth uniformly in the population. In contrast, two TnFLXgfp fusions in the *iolG* gene, encoding the inositol 2-dehydrogenase IolG, and the *spoVG* gene, encoding the RNA-binding regulatory protein SpoVG, gave rise to diffuse localization in the cytoplasm but only in a subset of cells in the population. Again, while lack of a subcellular localization pattern cannot be concluded with confidence based on loss-of-function insertion, the expression of the *iolG* and *spoVG* genes appears to be heterogeneous. We conclude that TnFLXgfp is useful for generating translational reporters that can also reveal subpopulation level gene expression in some cases.

Four TnFLXgfp insertion mutant strains showed fluorescence that was enriched at the cell periphery (Fig. 4C). Insertion of the transposon into the *mntB* gene, encoding the manganese transport protein MntB, exhibited fluorescence that was uniformly distributed throughout the periphery. The localization is consistent with the membrane and MntB is a membrane associated protein (35). Insertion of the transposon into three other loci including the gene *floT*, encoding the flotillin homolog FloT, the *fliG* gene, encoding the flagellar rotor protein FliG, and the *flhA* gene, encoding a component of flagellar type III export apparatus FlhA, exhibited a non-uniform or punctate localization at the membrane. We note that the localization pattern is consistent with previous observations in that FloT was reported to have a punctate localization (36), and FliG (37) and FlhA (38) are integral parts of the flagellar basal body that also localizes as puncta (39). We conclude that TnFLXgfp can randomly generate translational fusions within an open reading frame and can reproduce subcellular localization patterns that have been reported previously.

The last four TnFLXgfp strains indicated alternate localization patterns (Fig. 4D). For example, one mutant strain with an insertion into the coding region of the protease subunit ClpC, showed diffuse cytoplasmic localization with a sub fraction that accumulated as more intense puncta in the cytoplasm. The diffuse and punctate localization pattern of ClpC is consistent with some subcellular localization reports for this protein ((40)(41)). Two different insertions were found in the coding region of the chemotaxis protein CheV, and each insertion gave rise to intense cytoplasmic foci of variable size that sometimes appeared to be membrane-associated. Again, reports in *B. subtilis* and *Helicobacter pylori* support a punctate localization for CheV (42) (43). Finally, a strain harboring an insertion in the coding sequence of the alanine dehydrogenase SpoVN, produced filament-like structures that seemed to be localized near the cell periphery. Filamentous localization is an unusual pattern in bacteria, and while the localization of SpoVN in particular has not been directly investigated in *B. subtilis*, we note that other metabolic enzymes, e. g. the CTP synthase in the α-proteobacterium *C. crescentus*, has been reported to form filamentous structures ((44)(45)).

Here we generate a transposon delivery vehicle with increased stability in *E. coli* and that is amenable to easy modification. Moreover, we reproduce the previous-generation series of modified *mariner-based* transposons and develop a new reporter tool in the form a TnFLXgfp for making random fluorescent translational fusions *in vivo*. All plasmids will be donated to the Bacillus Genetic Stock Center (BGSC) for community acquisition. We note that while the new generation of plasmids are substantially more stable than the previous, propagation of transposons is inherently destabilizing due to spontaneous transposition, and thus we recommend minimal propagation and careful handling as a precaution.

## MATERIALS AND METHODS

### Strain construction

For the construction of the transposon delivery vectors, all plasmids were amplified in *E. coli* DH5α (see table 7 for all strains used in this study). To allow for transformation of *B. subtilis, E. coli* TG1 cells were transformed with the plasmids to generate concatenation. If not stated differently, all *B. subtilis* strains derived from the naturally competent modified undomesticated strain DK1042. Cells were incubated in lysogeny broth medium (5% (w/v) Yeast extract, 10% (w/v) tryptone, 0.09 M NaCl) or on plates containing this medium fortified with agar (1.5% (w/v)). Antibiotics were supplemented when appropriate at following concentrations: 100 μg/ml spectinomycin, 5 μg/ml kanamycin, 10 μg/ml tetracycline, 5 μg/ml chloramphenicol, mls (1 μg/ml erythromycin, 25 μg/ml lincomycin. For P_hyspank_ promoter dependent gene expression, 1 mM Isopropyl-β-d-1-thiogalactopyranoside (IPTG) was added to the medium. For colorimetric assays, using β-galactosidase activity as a readout, 0.5 mM X-Gal was supplemented.

**Table 5.** isolated TnFLXgfp mutants

| Strain number | Phenotype (localization) | Insertion | Tag | Annotation | Position of GFP insertion |
| --- | --- | --- | --- | --- | --- |
| DK7554 | Membrane | *mntB::TnFLXGFP cat* | TAAGGCTGA | manganese ABC transporter | MntB287 |
| DK7657 | Cytoplasm, heterogenous | *iolG::TnFLXGFP cat* | TATCAATCT | inositol 2-dehydrogenase | iolG336 |
| DK7658 | Cytoplasm and foci | *clpC::TnFLXGFP cat* | TAGAAGATG | ATPase subunit of the ClpC-ClpP protease | ClpC796 |
| DK7755 | Cytoplasm | *dps::TnFLXGFP kan* | TAGGTTAAG | iron storage protein | Dps144 |
| DK7552 | foci | *cheV::TnFLXGFP kan* | TAGAGGACT | Modulation of the two-component sensor kinase CheA | CheV194 |
| DK7662 | membrane | *fliG::TnFLXGFP kan* | TATCTTCTG | flagellar basal-body protein | FliG187 |
| DK7547 | cytoplasm | *fadR::TnFLXGFP tet* | TATCCAAAA | transcriptional repressor | FadR29 |
| DK7548 | foci | *cheV::TnFLX gfp tet* | TATCAAAAC | Modulation of the two-component sensor kinase CheA | CheV225 |
| DK7549 | membrane | *flhA::TnFLXGFP tet* | TAGATCCGA | part of the flagellar type III export apparatus | FlhA341 |
| DK7560 | cytoplasm | *spoVG::TnFLXGF P spec* | TAGATTCGA | RNA-binding regulatory protein | SpoVG89 |
| DK7559 | rods | *spoVN::TnFLXGF P spec* | TAGGCTATG | L-alanine dehydrogenase | SpoVN358 |
| DK7558 | membrane | *floT::TnFLXGFP spec* | TAATTGCAC | membrane-associated scaffold protein | FloT16 |

**Table 6.**
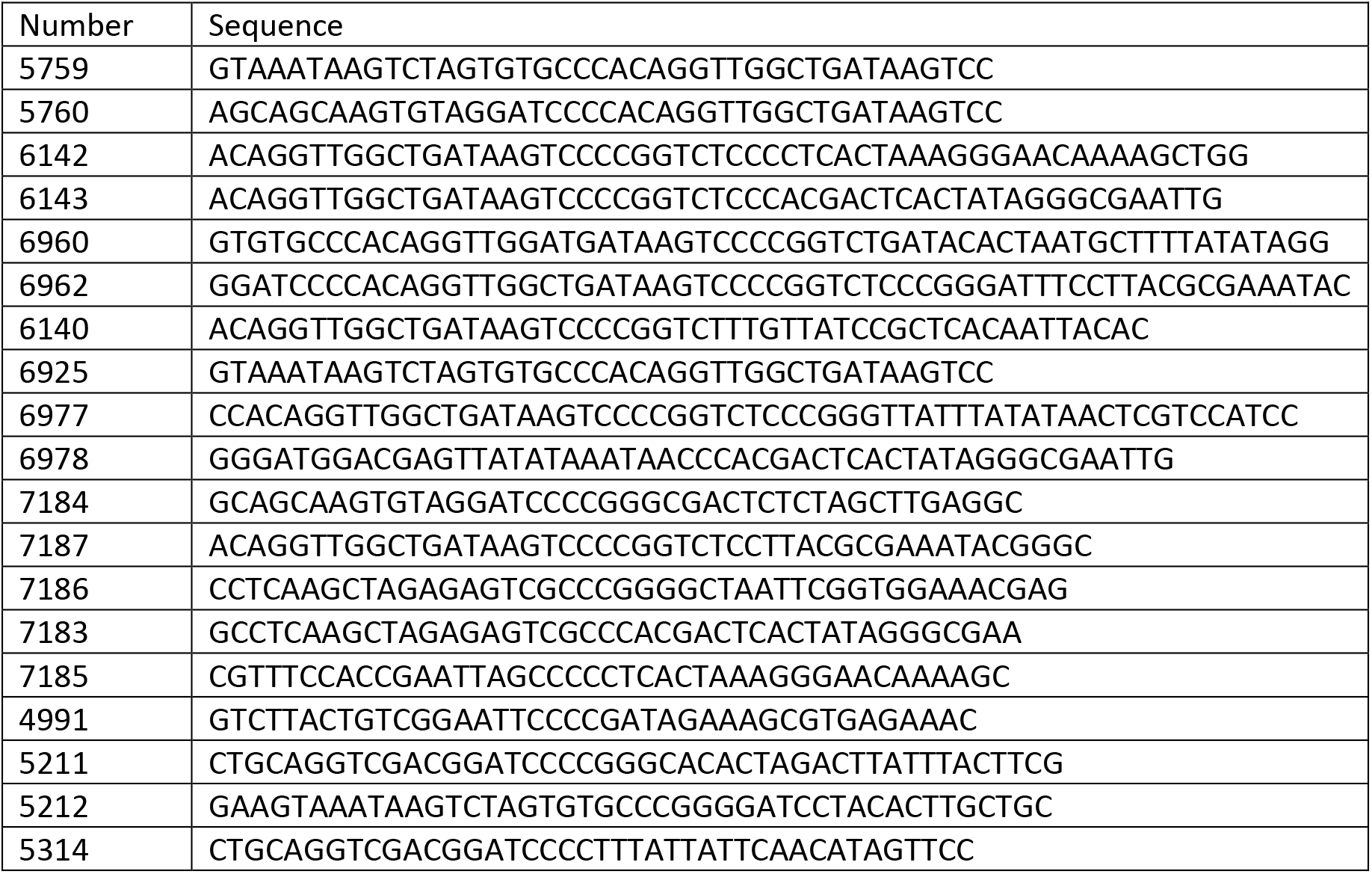
oligonucleotides used in this study

**Table 7.**
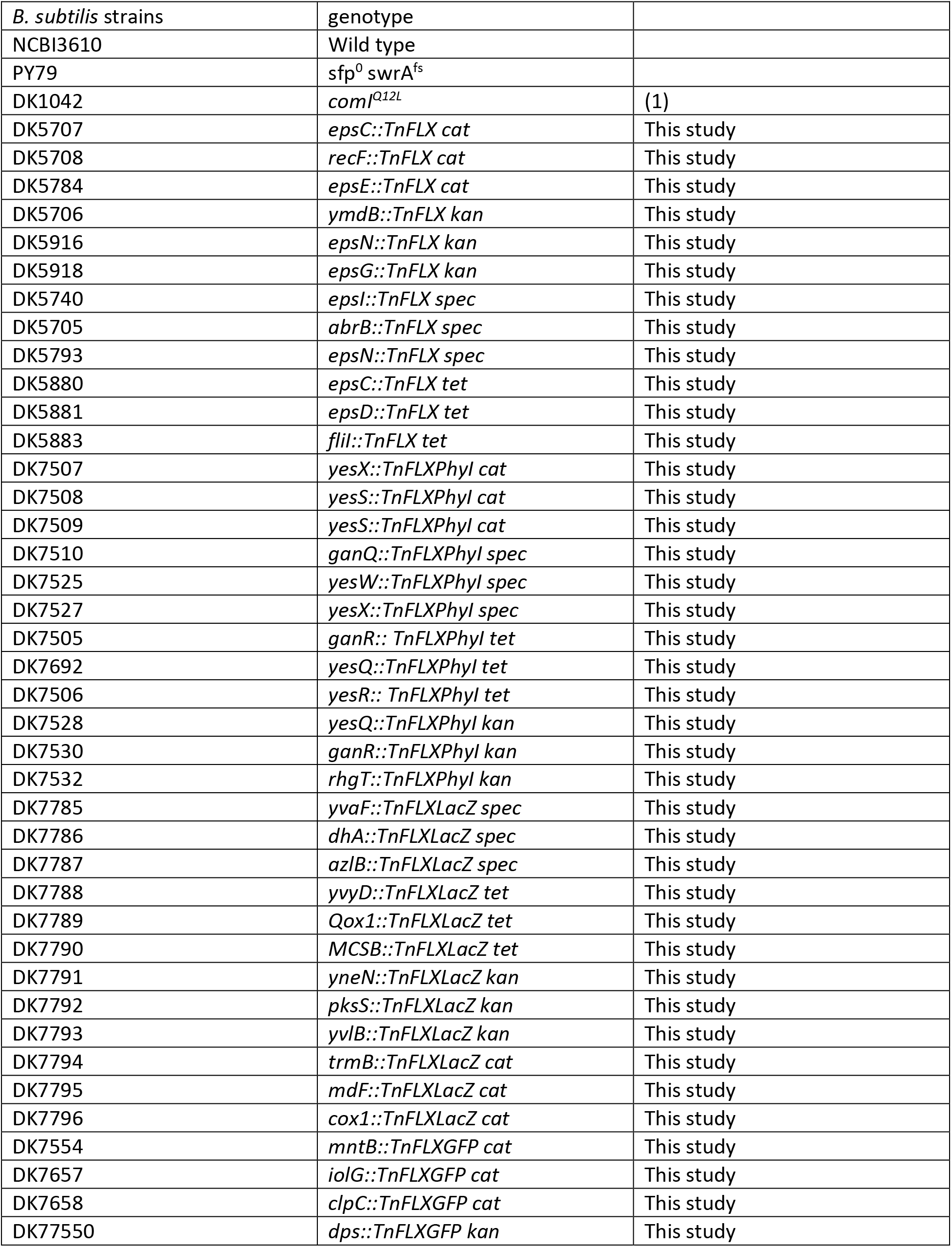
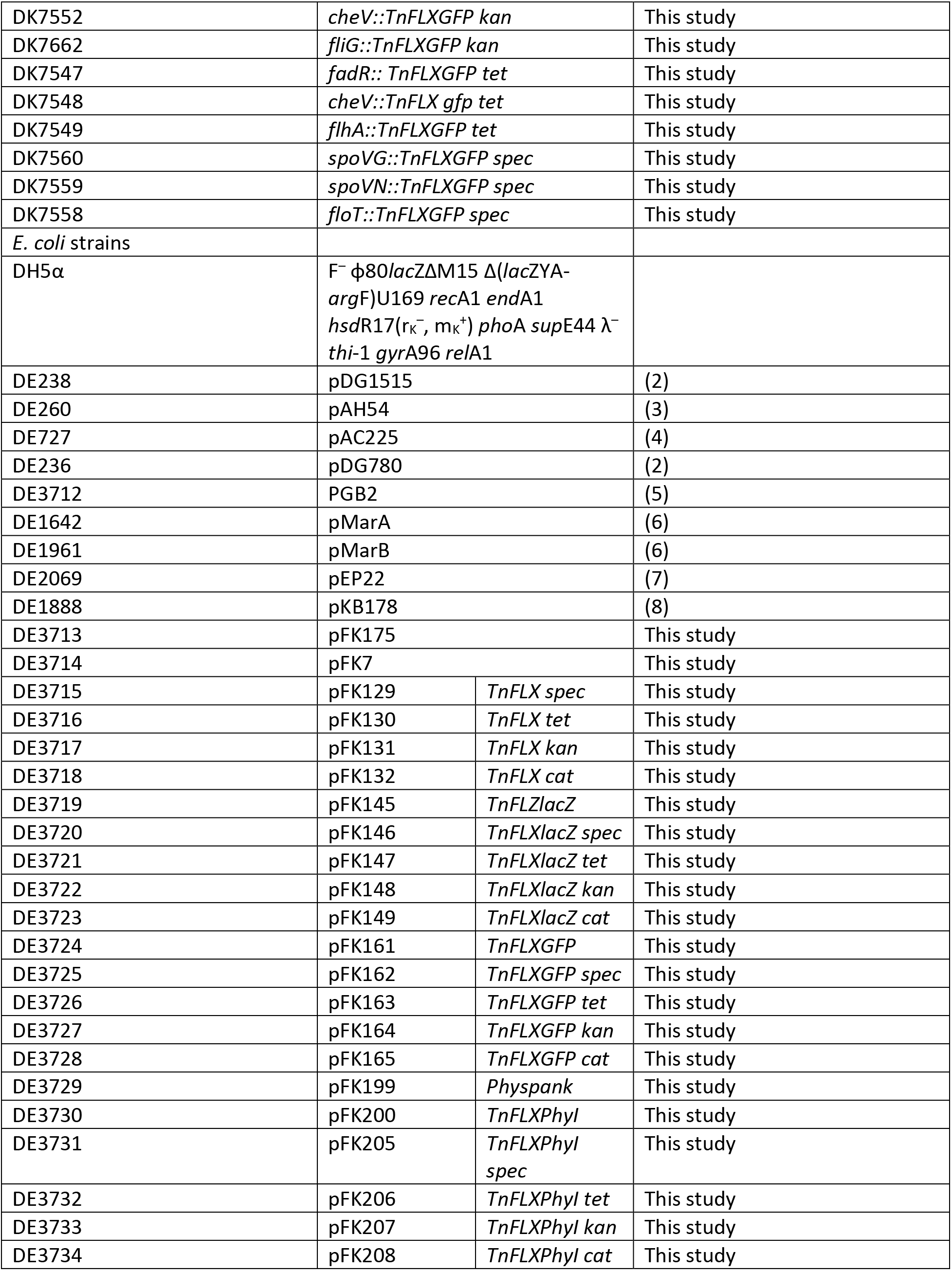
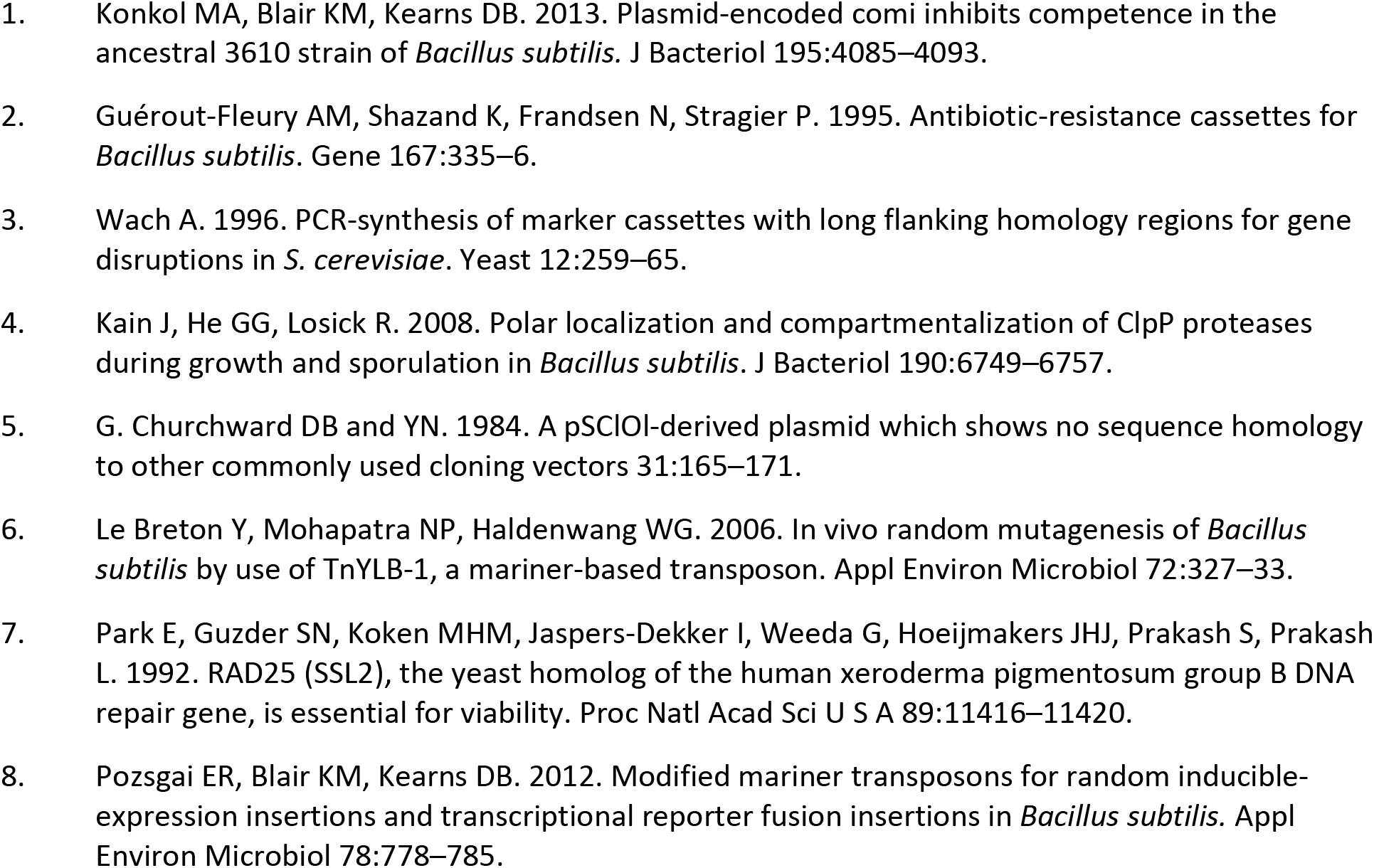
strains used in this study

### Plasmid construction

The plow copy-number plasmid pGB2 was linearized by *SmaI* treatment and complemented with an erythromycin resistance cassette (erm^R^) and a temperature sensitive origin of replication (oriBsTs), amplified from pMarB(2) using the oligonucleotides 5211 and 4991 by isothermal assembly (see table 6 for oligodesoxyribonucleotides used in this study), giving rise to pFK175. In a next step pFK175 was linearized with *SmaI* and the *mariner-Himar1* transposase coding region controlled by a σ^B^ dependent promoter, amplified from pMarB with the oligonucleotides 5212 and 5214, was integrated by isothermal assembly, creating pFK7. This plasmid was used as universal acceptor of all transposons described in this study.

The different transposons were constructed by PCR and integrated, as one or two DNA fragments respectively, into the *Sma*I linearized plasmid pFK7 by isothermal assembly.

### Transposons were constructed as follows

For all transposons the antibiotic resistance cassettes were amplified form from pDG1515 (46) (tetracycline resistance cassette (tet^R^)), pAH54 (47) (spectinomycin resistane cassette (spec^R^)), pAC225 (48)(chloramphenicol resistance cassette (cat^R^)), and pDG780 (46) (kanamycin resistance casstette (kan^R^)).

For the TnFLX series, the antibiotic resistance cassettes were amplified and extended with the inverted terminal repeats (ITRs) using the oligonucleotides 6142 and 6143. To allow for isothermal assembly, sequences overlapping with the *SmaI* linearized pFK7 plasmid were introduced by PCR with the oligonucleotides 5759 and 5760.

The TnFlexLacZ transposon series was assembled in a sequential manner. To Allow transcriptional coupling, first, the *lacZ* gene as well as a ribosomal binding site were amplified from pEP22 (49) and extended by ITR regions using the oligonucleotides 6960 and 6962, also integrating a *Sma*I recognition site. Homologies to the linearized pFK7 plasmid were introduced by an amplification step with the oligos 5759 and 5760. The resulting fragment was introduced into the *Sma*I linearized acceptor plasmid (pFK7) by isothermal assembly giving rise to pFK145. In a second step, antibiotic resistances cassettes were equipped with homologies to the acceptor plasmid by amplification with 6961 and 6142 and integrated into *Sma*I linearized pFK145 by ITA.

Transposons of the TnFLXPhyI series were assembled in three steps. First, the promoter encoding sequence was amplified from the plasmid pDP111 and extended by an ITR region by PCR with the oligonucleotides 6140 and 7184. The resulting fragment was equipped with homologies to *SmaI* linearized pFK7 by a second PCR using the primers 6925 and 7184 and integrated into the acceptor plasmid via ITA, creating pFK199. In a second step, the repressor encoging gene, *lacI*, was amplified from pDP111 and equipped with an ITR sequence using the primers 7186 and 7187. Regions overlapping with *Sma*I linearized pFK199 were introduced by PCR amplification with the oligos 7186 and 5760. Insertion of the resulting product into pFK199 was performed via ITA, giving rise to pFK200. The antibiotic resistance cassettes were introduced in a third step into *Sma*I linearized pFK200. Therefore, the coding regions were amplified and flanked by pFK200 homologies by PCR with the oligos 7183 and 7185 and integrated into pFK200 by ITA.

To generate a transposon allowing for the translational fusion of proteins with monomeric super folder green fluorescent protein (msfGFP), the coding sequence for *msfgfp* was analyzed, modified for optimal tRNA utilization according to the IDT codon optimization tool (www.idtdna.com) and synthesized by integrated DNA technologies (IDT). The obtained fragment, which already contained a single proximal ITR sequence, was equipped with a second, distal, ITR sequence by amplification with the oligos 6925 and 6977. To allow for integration into the acceptor plasmid, flanking homologies to pFK7 were introduced by PRC with 5759 and 5760, following an ITA reaction with *Sma*I linearized pFK7, creating pFK161. Antibiotic resistance cassettes were introduced into pFK161 by amplification of the coding region with the oligonucleotides 6978 and 5760 and a subsequent ITR reaction with *Sma*I linearized pFK161.

### Transposon mutagenesis

The transposon harboring plasmids were introduced into the *B. subtilis* acceptor strain via transformation (50), selecting for mls resistance and incubated at 30° C for 48 hours. Subsequently, cells were incubated for 16 hours at room temperature in LB-medium containing mls. The cell suspension was diluted 1:100 in LB medium containing the transposon specific antibiotic. The culture was grown for 6 hours at 42° C to cure the cells of the plasmid. To obtain individual colonies, the cell suspension was serially diluted and spread on LB plates, containing the appropriate antibiotic, and incubated overnight at 42° C.

### Transposon insertion site identification by inverse PCR (IPCR)

Cells were grown in LB medium at 37° for 6 hours. DNA was isolated and digested with the restriction endonuclease *Sau*3AI. Ligation with T4 DNA ligase was performed at room temperature for 2 hours. Inverse PCR was performed with 6 μl of ligated DNA using the oligonucleotides 6212 and 6420 (or 7195 and 7196 in the case of the TnFLXPhyi series) in a 50 μl reaction. The amplified fragment was analyzed by Sanger sequencing (IDT) using the universal sequencing oligo 6224.

### Fluorescence microscopy

Microscopy was performed with a Nikon eclipse 80i microscope, equipped with a Plan Apo 100X Ph 3 objective (NA 1.4). Cells were mounted on 1% (wt/vol) agarose pads containing S7_50_ minimal medium (51) on object slides. Images were acquired with a Cool snap HQ2 camera (Photometrix) and were processed with Metamorph 7.7.9 software (Universal Imaging Corp.). Membranes were stained with FM4-64 (final concentration, 1 nM; Molecular Probes).

### Flow Cytometry

Cells were grown in LB-medium at 37° C to an OD_600_ of 0.7, sedimented, and resuspended in phosphate buffered saline at a concentration of 5×10^6^ cells/ml. Analysis and cell sorting were performed, using a FACSAriaII flow cytometer (BD Biosciences) equipped with an 85 μm nozzle. Cells that were separated by FACS were collected in a reaction tubes containing LB medium that was complemented with the appropriate antibiotic and transferred onto fortified medium for further incubation.

## Acknowledgements

We thank Marta Perego for bringing to our attention the plasmid instability issue when plasmids carrying the TnKRM series was propagated in *E. coli*. We are grateful for the plasmids and the constructs received from the Bernhardt lab (Harvard university) and the Rudner lab (Harvard university). We thank Christiane Hassel (IUB flow cytometry core facility) for her support concerning bacterial cell sorting. This work was supported by NIH grant R35 GM131783 to DBK.

## Supplemental material

Sequence information of the relevant plasmids

